# Identification and expression of *Lactobacillus paracasei* genes for adaptation to desiccation and rehydration

**DOI:** 10.1101/475830

**Authors:** Aurore Palud, Karima Salem, Jean-François Cavin, Laurent Beney, Hélène Licandro

**Affiliations:** Université de Bourgogne Franche-Comté, AgroSup Dijon, PAM UMR A 02.102, Dijon, France; Ecole polytechnique de Sousse, Sousse, Tunisia

**Author notes:** Address correspondence to Aurore Palud, and Hélène Licandro.

**Keywords:** *Lactobacillus paracasei*, transposon mutants, desiccation, rehydration, gene expression

## Abstract

*Lactobacillus paracasei* is able to persist in a variety of natural and technological environments despite physico-chemical perturbations, in particular alternations between desiccation and rehydration. However, the way in which it adapts to hydric fluctuations and in particular the genetic determinants involved are not clearly understood. To identify the genes involved in adaptation to desiccation, an annotated library of *L. paracasei* random transposon mutants was screened for viability after desiccation (25% relative humidity, 25°C). Subsequently, the expression of the identified genes was measured at five stages of the dehydration-rehydration process to formulate the chronology of gene expression. The 24 identified genes were related to metabolism and transport, membrane function and structure, regulation of stress response, DNA related enzymes and environmental sensing. They were classified into four different transcriptomic profiles, in particular genes upregulated during both desiccation and rehydration phases and genes upregulated during the desiccation phase only. Thus, genetic response to hydric fluctuations seems to occur during desiccation and can continue or not during rehydration. The genes identified should contribute to improving the stabilization of lactobacillus starters in dry state.

**Importance:** Since water is the fundamental component of all living organisms, desiccation and rehydration alternation is one of the most prevalent and severe stresses for most microorganisms. Adaptation to this stress occurs via a combination of mechanisms which depend on the genetic background of the microorganism. In *L. paracasei,* we developed a strategy to identify genes involved in the adaptation to hydric fluctuations using random transposon mutagenesis and targeted transcriptomics. Both dehydration and rehydration were studied to decipher the chronology of genetic mechanisms. We found 24 as yet unidentified genes involved in this response. Most of them are linked to either the transport of molecules or to cell wall structure and function. Our screening also identified genes for environment sensing and two alarmones necessary for *L. paracasei* survival. Furthermore, our results show that desiccation is a critical phase for inducing stress response in *L. paracasei*.

## Introduction

Water is essential for all living organisms as it contributes to the structure of cells, stabilizes proteins, lipids and nucleic acids, and maintains vital metabolic systems and chemical reactions (1). Desiccation leads to water exit from cells, induces structural modifications, and causes osmotic and oxidative stresses (1–3). In addition, the rehydration phase could lead to membrane alterations (4). The capacity to return to life after hydric fluctuations is not only crucial for bacteria as a function of diurnal and seasonal cycles in natural environments but also during food industry processes such as conventional drying, freeze drying and spray drying (1, 5–7). In many organisms, desiccation survival is correlated with the accumulation of protective molecules, in particular trehalose (8). However, desiccation tolerance involves other mechanisms that are mostly unknown except for extreme desiccation tolerant bacteria (anhydrobiotes) such as such as cyanobacteria (9), yeast (2), resurrection plants (10) and microscopic animals (11).

In the last few years, progress has been made in understanding the mechanisms of bacteria tolerance to partial desiccation using transcriptome analysis, in particular for *Salmonella,* one of the most common foodborne pathogens able to survive for extended periods after desiccation (12–17). The comparison of these studies has revealed differences in identified genes that could be explained by the procedures used to dry bacteria (surface, drying medium, RH, desiccation periods). Several common genes were linked to amino acid transport and metabolism functions. Global transcriptional analyses were also performed for soil-residing bacteria (18–20) and similar responses were identified such as compatible solute and heat shock protein accumulation, reactive oxygen species neutralization, and DNA modification and repair. However, transcriptomics has limitations since genes essential for a function can be constitutively expressed and even inducible genes can be overexpressed during a very short period that does not overlap with that of the experiment. Recently, random transposon mutagenesis approaches were developed to identify genes for resistance to desiccation in *Listeria monocytogenes* (Hingston et al., 2015) and *Salmonella enterica* (22). For *L. monocytogenes* screening, genes were involved in energy production, membrane transport, amino acid metabolism, fatty acid metabolism and oxidative damage control. In *S. enterica*, in comparison to previous transcriptomic studies, several genes were related to amino acid metabolism and more than 20% encoded hypothetical proteins. Recently, changes in proteomic expression were investigated for *S. enterica* during desiccation and rehydration (23). The proteins with higher expression levels in dried samples were mainly ribosomal proteins whereas flagellar proteins, membrane proteins, and export systems as well as stress response proteins were identified in rehydrated samples.

Although *L. casei/paracasei* is one of the most emblematic groups of lactic acid bacteria (LAB), the genetic mechanisms involved in desiccation resistance and adaptation are not fully understood, limiting prospects for improving and developing preservation processes. Comparative genomics has demonstrated that LAB are highly adaptable to various niches such as soil, foods (dairy, meat, and vegetable), as well as oral, vaginal and gastrointestinal cavities (24–26). This is correlated with an ability to adapt and persist under diverse environmental stresses (27). Thanks to this capacity, humans have succeeded in selecting and producing certain LAB strains as efficient starters for fermented foods (particularly hard cheese) and as probiotics. The adequate preservation of starter cultures is necessary though it is currently only possible using advanced desiccation technologies and storage in frozen state (28)

Considering the importance of the *L.casei/paracasei* group for health and industrial applications, we compiled a non-redundant, annotated transposon mutant library of *L. paracasei* (29, 30) based on the P_junc_-TpaseIS_*1223*_ system, a random mutagenesis tool specifically designed for the *Lactobacillus* genus (31). Recently, using this approach we determined five *Lactobacillus pentosus* mutants sensitive to olive brine due to multifactorial stress (32). In addition, new genetic determinants were identified during the early stage of *L. paracasei* establishment in the gut (29) and during monofactorial perturbations of mild intensities (33). In the present work, the *L. paracasei* transposon mutant library was screened to identify genes involved in the adaptation of this LAB to desiccation and rehydration. The expression of corresponding genes was studied by RT-qPCR during the two-step process to draw a chronological transcriptomic profile during hydric fluctuations.

## Results

### Development of the screening strategy

A library containing a total number of 1287 *L. paracasei* mutants was screened for their capacity to resist a 24-h desiccation period followed by rapid rehydration in comparison with their parental strain (*L. paracasei* ATCC 334, named WT). Convective drying in a ventilated chamber at room temperature with a relative humidity (RH) of 25% was selected to mimic the severe desiccation conditions that can occur in natural environments during drought in soil, on plants, and on animal skin, for example. This process consisted in passing a flow of dry air over the cell suspension approximately simulating the desiccation conditions that organisms can undergo in the natural environment. To assess the importance of sugars as drying protectors, we selected three disaccharides related in the literature to lactic acid bacteria preservation, in particular during freeze drying (34–36), and their corresponding monosaccharides. The viability of *L. paracasei* WT after 24-h desiccation at 25% RH was determined with and without the protectors (Table 1). Viability was very low in the absence of sugar and with monosaccharides. On the contrary, with disaccharides, viability was between 19 and 45%, which was an adequate range for screening either less or more resistant mutants. Lactose 50 g/L was selected because it corresponds to the composition of milk, thus a condition often encountered by *L. paracasei* in the natural and technological environments of cheese-making, for example. The whole strategy followed in this work is illustrated in figure 1. A screening method composed of easy and rapid steps to save time and reduce the risk of bias was developed. The viability of the 1287 mutants after 24-h desiccation followed by rapid rehydration was evaluated in 96-well microplates by monitoring the acidification kinetics (due to sugar metabolism) using a medium containing a pH indicator. Absorbance (A_420nm_) was correlated to the quantity of metabolically active (or viable) bacteria, thus the viability of the mutants could be compared to that of the WT (A_420nm_ (7 h) = 1.91 ± 0.08). Sensitive and resistant mutants were subsequently validated by individual drying on PP coupons and by determining viability by the plate count method in order to remove possible false positive mutants (for instance, metabolic deficiency mutants).

**Table 1.**
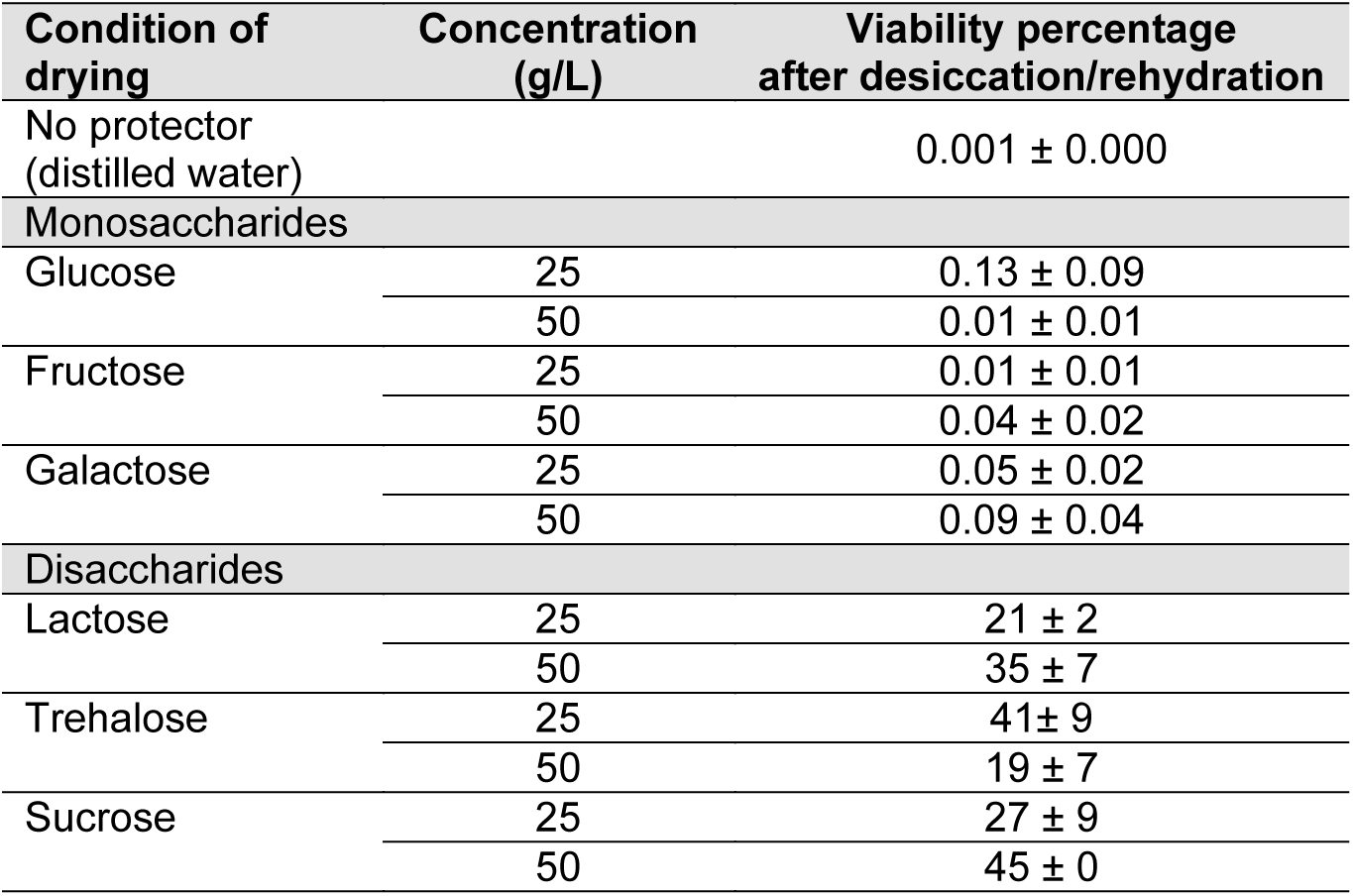
*L. paracasei* ATCC 334 viability after a pronounced desiccation period followed by rapid rehydration with or without saccharides used as protectors. Bacterial cells in stationary growth phase suspended in various protective solutions were air dried for 24 h at 25% RH and 25°C on PP coupons. Survival was determined after suspending dried cells in BCP medium. Viability was measured by plate count (CFU/ml).

| Condition of drying | Concentration (g/L) | Viability percentage after desiccation/rehydration |
| --- | --- | --- |
| No protector (distilled water) |  | 0.001 ± 0.000 |
| **Monosaccharides** |  |  |
| Glucose | 25 | 0.13 ± 0.09 |
|  | 50 | 0.01 ± 0.01 |
| Fructose | 25 | 0.01 ± 0.01 |
|  | 50 | 0.04 ± 0.02 |
| Galactose | 25 | 0.05 ± 0.02 |
|  | 50 | 0.09 ± 0.04 |
| **Disaccharides** |  |  |
| Lactose | 25 | 21 ± 2 |
|  | 50 | 35 ± 7 |
| Trehalose | 25 | 41 ± 9 |
|  | 50 | 19 ± 7 |
| Sucrose | 25 | 27 ± 9 |
|  | 50 | 45 ± 0 |

**Figure 1.**
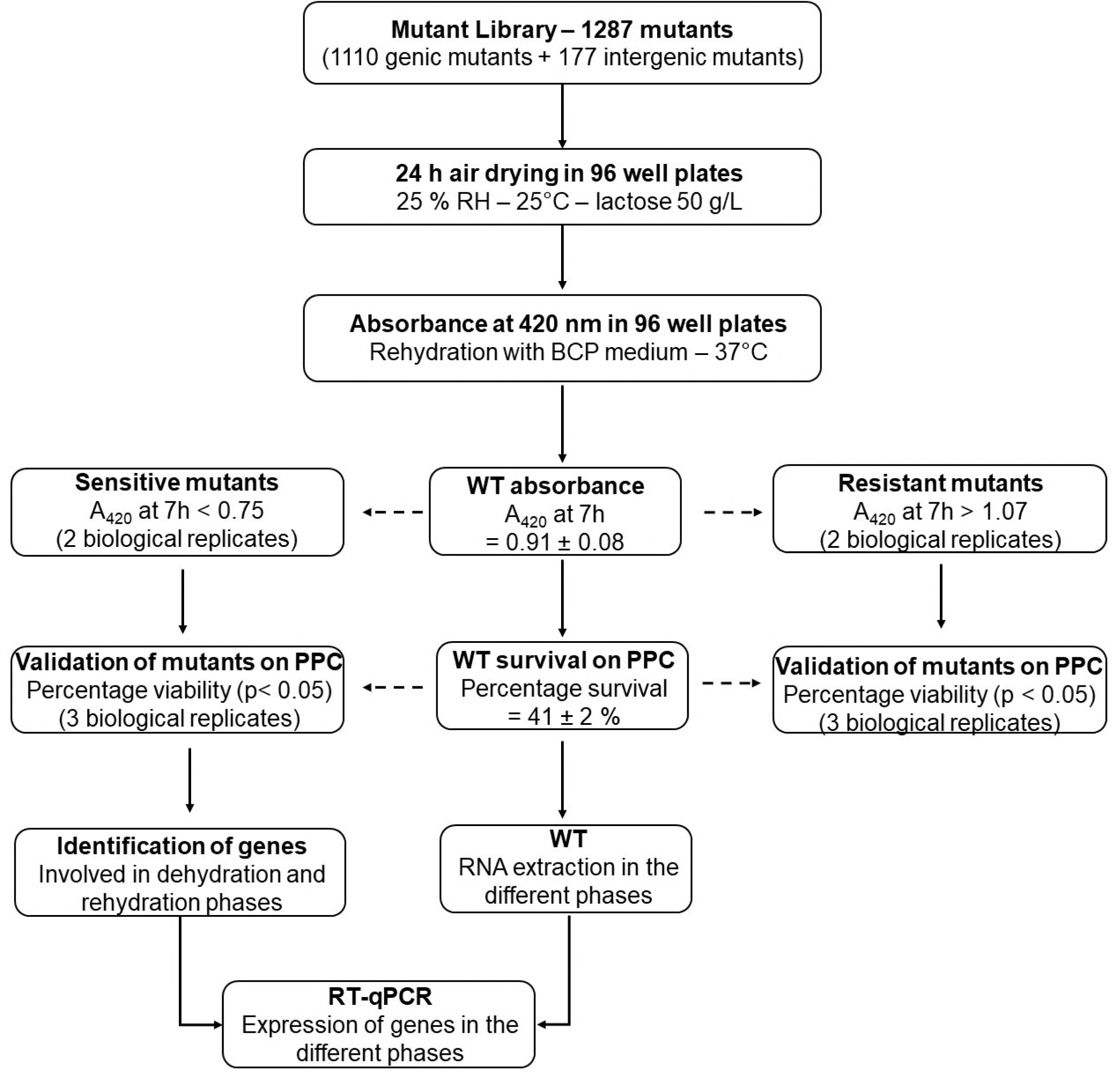
Organizational chart of the strategy developed in this work to identify genetic determinants of *L. paracasei* resistance to desiccation and rehydration. PPC, polypropylene coupons.

### Identification of genes involved in resistance to pronounced desiccation followed by rapid rehydration

The mutant library was subjected to 24-h desiccation (pronounced desiccation) followed by rapid rehydration. Screening of the 1287 transposon mutants resulted in the detection of 47 sensitive mutants (A_420nm_ at 7 h < 1.75 for 2 biological replicates) but no resistant mutants (A_420nm_ at 7 h > 2.07 for 2 biological replicates). After the validation step by individual drying of the 47 mutants, 24 mutants displayed a significant decrease of survival compared to the WT (p<0.05) (Table 2). Identified genes were mostly involved in transport, regulation and membrane binding proteins. Moreover, five sensitive mutants containing transposon in a gene encoding a hypothetical protein were identified (LSEI_0040, LSEI_0733, LSEI_0806, LSEI_1045 and LSEI_1316). Mutants were classified into three categories according to their viability: + (mean viability at least 1.2-fold less), ++ (mean viability at least 2-fold less) and +++ (mean viability at least 3-fold less than WT). The majority of mutants (20) were categorized as +. The two most sensitive mutants (+++) were disrupted for a putative ribonucleotide diphosphate reductase (LSEI_1468) and for a hypothetical protein (LSEI_ 1316). Two mutants disrupted for a putative PTS (LSEI_0178) and for a putative alpha/beta fold family hydrolase (LSEI_0756) were categorized as ++. Fifteen genes out of the 24 belonged to putative operons (Table 3). Eleven genes were specific to *L. paracasei* or related species, five to *Lactobacillus* genus and eight were well conserved among Gram (+) bacteria. Two genes among the five encoding hypothetical proteins presented a transmembrane domain (LSEI_0040 and LSEI_1045). All hypothetical protein genes, except LSEI_0733, were specific to the *L. paracasei* group.

**Table 2.** *L. paracasei* mutants sensitive to a pronounced desiccation period followed by rapid rehydration and their corresponding A_420nm_ at 7h and viability percentage. Cells in stationary growth phase were air dried for 24 h at 25% RH and 25°C with 50 g/L lactose solution in microplates or on PP coupons. Survival was determined by monitoring A420nm of dried cells suspended in BCP medium. Mutants were classified into three categories according to their viability: + (mean viability at least 1.2-fold less), ++ (mean viability at least 2-fold less) and +++ (mean viability at least 3-fold less than WT.

| Disrupted gene | Predictive function | $A_{420nm}$ at 7 h<br>2 repetitions | Viability %<br>3 repetitions | Sensitivity level |
| --- | --- | --- | --- | --- |
| WT | | $1.91 \pm 0.08$ | $41 \pm 2$ | |
| LSEI_0040 | Hypothetical protein | 1.56 / 1.69 | $23 \pm 4$ | + |
| LSEI_0167 | Diadenosine tetraphosphatase | 1.71 / 1.64 | $23 \pm 3$ | + |
| LSEI_0178 | PTS | 1.73 / 1.59 | $18 \pm 9$ | ++ |
| LSEI_0238 | PST transporter | 1.63 / 1.66 | $23 \pm 7$ | + |
| LSEI_0363 | Pyridoxine 5'-phosphate oxidase | 1.74 / 1.75 | $26 \pm 6$ | + |
| LSEI_0397 | NAD (FAD)-dependent dehydrogenase | 1.34 / 1.40 | $25 \pm 5$ | + |
| LSEI_0733 | Hypothetical protein | 1.74 / 1.75 | $31 \pm 5$ | + |
| LSEI_0749 | Cation transport ATPase | 1.74 / 1.75 | $23 \pm 3$ | + |
| LSEI_0756 | Alpha/beta fold family hydrolase | 1.62 / 1.75 | $18 \pm 1$ | ++ |
| LSEI_0758 | Aldo/keto reductase related enzyme | 1.45 / 1.48 | $22 \pm 3$ | + |
| LSEI_0806 | Hypothetical protein | 1.46 / 1.59 | $29 \pm 3$ | + |
| LSEI_0934 | DNA-binding response regulator | 1.48 / 1.59 | $28 \pm 4$ | + |
| LSEI_1009 | Spermidine/putrescine-binding protein | 1.63 / 1.45 | $26 \pm 1$ | + |
| LSEI_1045 | Hypothetical protein | 1.72 / 1.69 | $23 \pm 4$ | + |
| LSEI_1316 | Hypothetical protein | 1.30 / 1.27 | $14 \pm 6$ | +++ |
| LSEI_1324 | Membrane metal-binding protein | 1.70 / 1.63 | $33 \pm 3$ | + |
| LSEI_1468 | Ribonucleotide diphosphate reductase | 1.10 / 1.11 | $4 \pm 1$ | +++ |
| LSEI_1539 | ppGpp | 1.35 / 1.56 | $29 \pm 5$ | + |
| LSEI_1659 | Glucokinase | 1.65 / 1.66 | $25 \pm 8$ | + |
| LSEI_1709 | DNA polymerase I | 1.65 / 1.60 | $28 \pm 6$ | + |
| LSEI_1754 | SAICAR synthase | 1.75 / 1.46 | $23 \pm 4$ | + |
| LSEI_2505 | Phosphate-starvation-inducible protein | 1.71 / 1.75 | $32 \pm 3$ | + |
| LSEI_2725 | Sorbitol transcription regulator | 1.74 / 1.70 | $27 \pm 5$ | + |
| LSEI_A05 | PTS cellobiose-specific | 1.59 / 1.44 | $31 \pm 2$ | + |

**Table 3.** Operon organization and specificity of the 24 genes involved in adaptation to pronounced desiccation followed by rapid rehydration.

| Gene | Predictive function | Putative operon | DNA strand | Specificity |
| --- | --- | --- | --- | --- |
| LSEI_0040 | Hypothetical protein | No | (+) | *L.casei/paracasei/rhamnosus* |
| LSEI_0167 | Diadenosine tetraphosphatase | No | (-) | *L.casei/paracasei/rhamnosus* |
| LSEI_0178 | PTS | 0178-0180 | (+) | *L.casei/paracasei/rhamnosus* |
| LSEI_0238 | PST transporter | 0238-0240 | (+) | *L.casei/paracasei/rhamnosus* |
| LSEI_0363 | Pyridoxine 5'-phosphate oxidase | No | (-) | *Lactobacillus* |
| LSEI_0397 | NAD (FAD)- dehydrogenase | No | (-) | *Lactobacillus* |
| LSEI_0733 | Hypothetical protein | 0731-0733 | (-) | Gram + |
| LSEI_0749 | Cation transport ATPase | No | (-) | *Lactobacillus* |
| LSEI_0756 | Alpha/beta fold family hydrolase | No | (+) | *Lactobacillus* |
| LSEI_0758 | Aldo/keto reductase related enzyme | 0757-0758 | (-) | Gram + |
| LSEI_0806 | Hypothetical protein | 0805-0806 | (+) | *L.casei/paracasei/rhamnosus* |
| LSEI_0934 | DNA-binding response regulator | 0933-0941 | (+) | *Lactobacillus* |
| LSEI_1009 | Spermidine/putrescine-binding protein | 1005-1009 | (+) | Gram + |
| LSEI_1045 | Hypothetical protein | No | (-) | *L.casei/paracasei/rhamnosus* |
| LSEI_1316 | Hypothetical protein | 1314-1319 | (+) | *L.casei/paracasei/rhamnosus* |
| LSEI_1324 | Membrane metal-binding protein | 1320-1326 | (+) | *L.casei/paracasei/rhamnosus* |
| LSEI_1468 | Ribonucleotide reductase | 1467-1470 | (+) | Gram + |
| LSEI_1539 | ppGpp synthase | 1537-1539 | (-) | Gram + |
| LSEI_1659 | Glucokinase | 1658-1661 | (-) | Gram + |
| LSEI_1709 | DNA polymerase | 1704-1709 | (-) | Gram + |
| LSEI_1754 | SAICAR synthase | 1746-1756 | (-) | Gram + |
| LSEI_2505 | Phosphate-starvation-inducible protein | No | (+) | *L.casei/paracasei/rhamnosus* |
| LSEI_2725 | Sorbitol transcription regulator | 2720-2726 | (-) | *L.casei/paracasei/rhamnosus* |
| LSEI_A05 | PTS | A04-A06 | (-) | *L.casei/paracasei/rhamnosus* |

### Analysis of mutant sensitivity after pronounced desiccation followed by progressive rehydration

To assess if the identified genes were involved during dehydration whatever the rehydration process, the WT and the 24 sensitive mutants identified previously were subjected to desiccation for 24-h followed by progressive rehydration (for 2 h) instead of rapid rehydration. In these conditions, the viability of the WT was 99 ± 3%, which is considerably higher than obtained with a rapid rehydration. Among the 24 sensitive mutants identified previously, only seven were also sensitive after desiccation and progressive rehydration (Table 4). They encode a putative diadenosine tetraphosphatase (LSEI_0167), a PTS system (LSEI_0178), an NAD (FAD)-dependent dehydrogenase (LSEI_0397), a cation transport ATPase (LSEI_0749), a ribonucleotide reductase (LSEI_1468), a SAICAR synthase (LSEI_1754) and a hypothetical protein (LSEI_0040). As observed for desiccation and rapid rehydration, the most sensitive mutant was ribonucleotide diphosphate reductase.

**Table 4.** *L. paracasei* mutants sensitive to pronounced desiccation followed by progressive rehydration and their corresponding viability percentage among mutants identified as sensitive to pronounced desiccation followed by rapid rehydration. Cells in stationary growth phase were air dried for 24 h at 25% RH and 25°C with 50 g/L lactose solution on PP coupons. Progressive rehydration was performed in a closed chamber with RH adjusted at 99% for 2 h and the addition of water to obtain the mass before drying. Viability percentages were determined by suspending dried cells with BCP medium.

| Disrupted gene | Predictive function | Viability %<br>3 repetitions |
| --- | --- | --- |
| **WT** |  | **99 ± 3** |
| LSEI_0040 | Hypothetical protein | 86 ± 7 |
| LSEI_0167 | Diadenosine tetraphosphatase | 60 ± 0 |
| LSEI_0178 | PTS | 82 ± 7 |
| LSEI_0397 | NAD (FAD)-dependent dehydrogenase | 41 ± 6 |
| LSEI_0749 | Cation transport ATPase | 57 ± 4 |
| LSEI_1468 | Ribonucleotide diphosphate reductase | 14 ± 2 |
| LSEI_1754 | SAICAR synthase | 87 ± 8 |

### Transcriptomic analysis of the identified genes during a desiccation – rehydration cycle

Mutant library screening allowed the identification of 24 genes involved in the global perturbation which consisted in desiccation followed by rehydration. Thus, it was not possible to determine during which phase these genes were involved since rehydration was necessary to analyze mutant phenotypes. To determine the chronology of the genetic mechanisms during humidity fluctuations, transcriptomic analysis of the identified genes was carried out on the WT. RNA extractions were performed at two different stages of desiccation: partial desiccation D1 (76% of water evaporated after 1 h) and pronounced desiccation D2 (96% of water evaporated after 2 h) which corresponded to completely dried cells (Figure 2). As rehydration kinetics is decisive for bacterial survival, rapid and progressive rehydration were applied for RNA extractions after 2 h.

**Figure 2.**
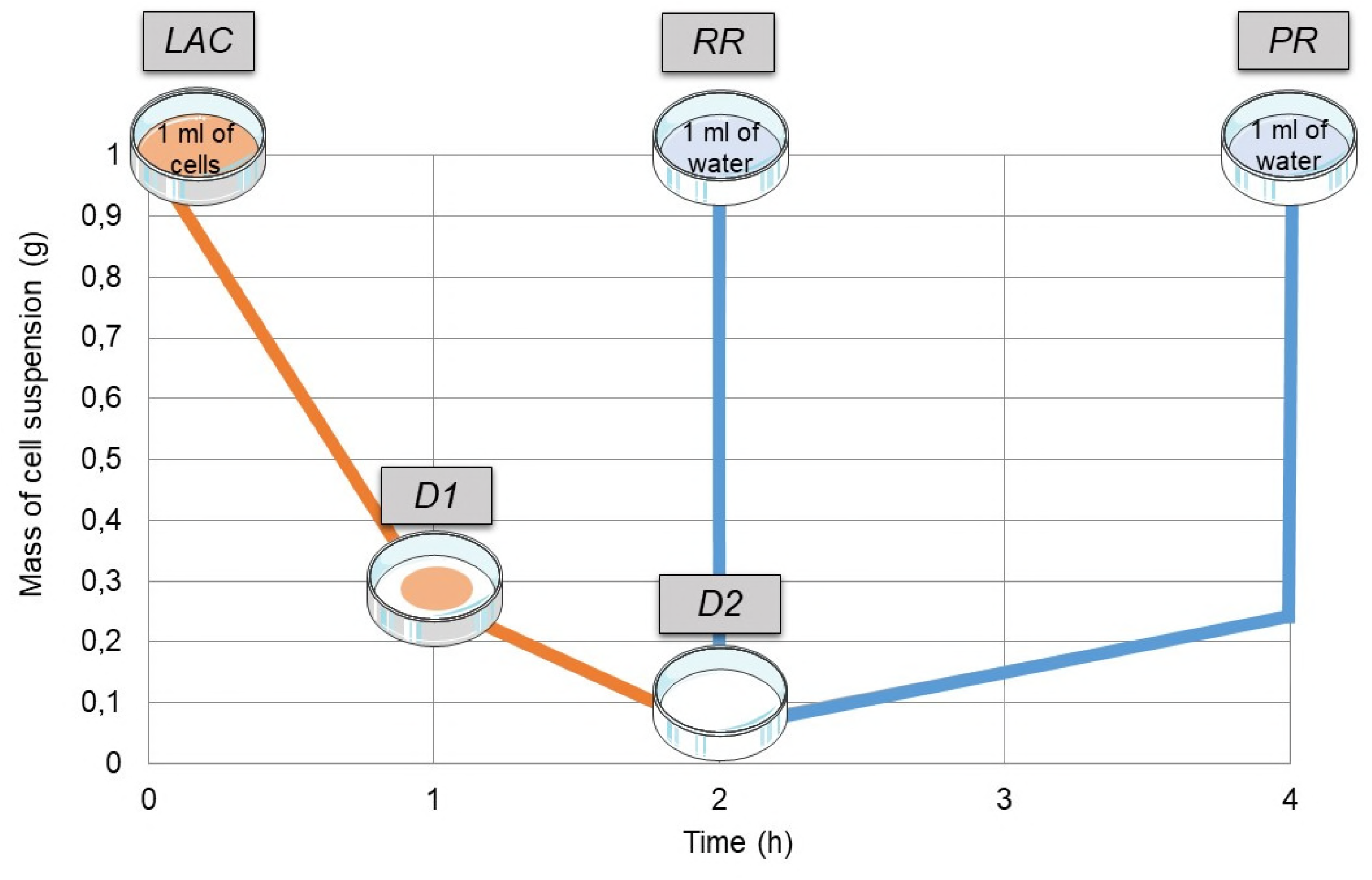
Evolution of the cell suspension mass during desiccation (orange curve) and rehydration (blue curve) on a polyvinylidene membrane. Sampling times for RNA extractions are represented by the grey boxes: LAC (1 mL of cells incubated 15 min with lactose before drying), D1 (cells dried 1 h, partial desiccation), D2 (cells dried 2 h, pronounced desiccation), RR (rapid rehydration with 1 mL of distillated water), PR (progressive rehydration in a closed chamber with RH adjusted at 99% for 2 h and addition of water to obtain the mass before drying).

As lactose was required to ensure good survival of bacteria during the desiccation/rehydration process it was reasonable to expect that some candidate genes could be differentially expressed in the presence of lactose. However, for the 24 genes, expressions after 15-min incubation in lactose were comparable to that after a 15-min phosphate buffer incubation (Figure 3). Differentially expressed genes were classified to obtain the most upregulated (Table 5, values in dark red) and downregulated (Table 5, values in dark blue) genes, with a mean expression value higher or lower than 2.0. Among the 24 genes studied, 18 were differentially expressed (p < 0.05) for at least one condition (Table 5). The six genes involved in hydric changes but not differentially expressed encoded a putative PTS transporter (LSEI_0178), a reductase (LSEI_0758), a ppGpp alormone (LSEI_1539), a DNA polymerase (LSEI_1709) and hypothetical proteins (LSEI_0733 and LSEI_1045). Six genes were upregulated for all the hydric fluctuations tested. These genes encoded a putative diadenosine tetraphosphatase (LSEI_0167), a polysaccharide transporter (LSEI_0238), a pyridoxine 5’- phosphate oxidase (LSEI_0363), a PTS transporter (LSEI_A05) and hypothetical proteins (LSEI_0040 and LSEI_0806). Also, LSEI_0397 encoding a putative NAD dependent dehydrogenase was upregulated during desiccation and progressive rehydration. Conversely, LSEI_0749 encoding a putative cation transport ATPase was upregulated during desiccation and rapid rehydration. Interestingly, these genes were the most upregulated (> 2-fold change) for at least one condition. LSEI_A05 (putative lactose specific PTS system) was the most upregulated gene during desiccation (D1 and D2). One gene, LSEI_1754, encoding a putative SAICAR synthase was the most downregulated gene in these experiments (mean expression value < 2.0 for all the drying and rehydration conditions).

**Table 5.** Relative transcript level of *L paracasei* genes after desiccation (1 h and 2 h) and rehydration (rapid or progressive) for the 24 genes identified after library mutant screening. LAC (cells incubated 15 min with lactose), D1 (cells dehydrated for 1 h), D2 (cells dehydrated for 2 h), RR (rapid rehydration with 1 mL of distillated water), PR (progressive rehydration in a closed chamber with RH adjusted at 99% for 2 h). RTL were calculated using 2^-ΔΔCt^ method. For the phosphate buffer control condition, a gene expression value of 1.0 was attributed and RTL of genes in stress condition were calculated as a function of this value. Positive values (> 1.0) represent upregulation and negative values (< 1.0) represent downregulation. *, significant changes in gene expression (p<0.05) compared to the phosphate buffer condition (4 biological replicates). Values in light red correspond to upregulation and in dark red to upregulations > 2.0; values in light blue correspond to downregulation and in dark blue to down regulation < −2.0.

| Gene | Fonction | LAC | D1 | D2 | RR | PR |
| --- | --- | --- | --- | --- | --- | --- |
| LSEI_0040 | Hypothetical protein | 1.0 ± 0.1 | 1.7 ± 0.6* | 2.6 ± 0.3* | 2.1 ± 0.2* | 1.9 ± 0.4 |
| LSEI_0167 | Diadenosine tetraphosphatase | 1.0 ± 0.1 | 1.8 ± 0.4* | 2.3 ± 0.3* | 1.6 ± 0.2 | 2.1 ± 0.4* |
| LSEI_0178 | PTS | 1.0 ± 0.2 | -1.1 ± 0.2 | -1.1 ± 0.1 | -1.3 ± 0.2 | -1.3 ± 0.3 |
| LSEI_0238 | PST family polysaccharide transporter | 1.0 ± 0.1 | 1.3 ± 0.5* | 2.2 ± 0.3* | 1.6 ± 0.1* | 1.9 ± 0.3* |
| LSEI_0363 | Pyridoxine 5'-phosphate oxidase | 1.0 ± 0.1 | 1.4 ± 0.4* | 1.9 ± 0.2* | 1.6 ± 0.2* | 1.9 ± 0.3* |
| LSEI_0397 | NAD(FAD)-dependent dehydrogenase | 1.0 ± 0.2 | 1.6 ± 0.4* | 1.8 ± 0.2* | 1.7 ± 0.2* | -1.3 ± 0.2 |
| LSEI_0733 | Hypothetical protein | 1.1 ± 0.1 | -1.1 ± 0.2 | 1.3 ± 0.2 | 1.1 ± 0.1 | 1.2 ± 0.3 |
| LSEI_0749 | Cation transport ATPase | 1.1 ± 0.1 | 2.1 ± 0.2* | 2.1 ± 0.2* | 1.3 ± 0.3 | 1.6 ± 0.1* |
| LSEI_0756 | Alpha/beta fold family hydrolase | 1.0 ± 0.2 | 1.2 ± 0.1 | 1.1 ± 0.1 | -1.4 ± 0.2* | -1.7 ± 0.3* |
| LSEI_0758 | Aldo/keto reductase | -1.1 ± 0.2 | -1.4 ± 0.4 | -1.1 ± 0.1 | -1.1 ± 0.1 | 1.0 ± 0.2 |
| LSEI_0806 | Hypothetical protein | -1.1 ± 0.1 | 1.7 ± 0.5* | 2.3 ± 0.3* | 2.0 ± 0.2* | 2.4 ± 0.4* |
| LSEI_0934 | DNA-binding response regulator | -1.1 ± 0.1 | 1.2 ± 0.2 | 1.6 ± 0.2* | 1.2 ± 0.1 | 1.4 ± 0.2 |
| LSEI_1009 | Spermidine/putrescine-binding protein | -1.1 ± 0.1 | -1.3 ± 0.2 | -1.4 ± 0.1 | -2.0 ± 0.4* | -2.0 ± 0.4* |
| LSEI_1045 | Hypothetical protein | -1.1 ± 0.1 | 1.2 ± 0.1 | 1.2 ± 0.2 | -1.3 ± 0.2 | -1.4 ± 0.2 |
| LSEI_1316 | Hypothetical protein | -1.1 ± 0.1 | 1.7 ± 0.2* | 1.6 ± 0.2* | 1.0 ± 0.1 | -1.1 ± 0.2 |
| LSEI_1324 | Membrane metal-binding protein | -1.1 ± 0.1 | -1.4 ± 0.2 | -1.3 ± 0.1 | -1.4 ± 0.2* | -1.3 ± 0.2 |
| LSEI_1468 | Ribonucleotide diphosphate reductase | 1.0 ± 0.2 | 1.5 ± 0.1 | 1.2 ± 0.1 | -1.3 ± 0.2* | -1.7 ± 0.3* |
| LSEI_1539 | ppGpp synthase | 1.0 ± 0.1 | -1.3 ± 0.2 | -1.1 ± 0.1 | -1.1 ± 0.1 | -1.3 ± 0.3 |
| LSEI_1659 | Glucokinase | 1.1 ± 0.1 | -1.4 ± 0.4 | -1.3 ± 0.2 | -1.4 ± 0.2* | -1.4 ± 0.2 |
| LSEI_1709 | DNA polymerase | 1.0 ± 0.2 | 1.0 ± 0.3 | 1.2 ± 0.2 | 1.2 ± 0.1 | 1.3 ± 0.3 |
| LSEI_1754 | SAICAR synthase | -1.1 ± 0.1 | -2.5 ± 0.6* | -2.5 ± 0.6* | -2.5 ± 0.6* | -2.5 ± 0.6* |
| LSEI_2505 | Phosphate-starvation-inducible protein | -1.1 ± 0.1 | -1.1 ± 0.4 | 1.8 ± 0.2* | 1.2 ± 0.2 | 1.1 ± 0.3 |
| LSEI_2725 | Sorbitol operon transcription regulator | -1.1 ± 0.1 | 1.3 ± 0.3 | 1.5 ± 0.2* | 1.2 ± 0.1 | 1.4 ± 0.3 |
| LSEI_A05 | PTS | 1.0 ± 0.1 | 3.5 ± 0.2* | 4.1 ± 0.2* | 2.3 ± 0.1* | 2.1 ± 0.3* |
| **Total of genes differentially expressed** |  | **0** | **10** | **13** | **13** | **11** |

**Figure 3.**
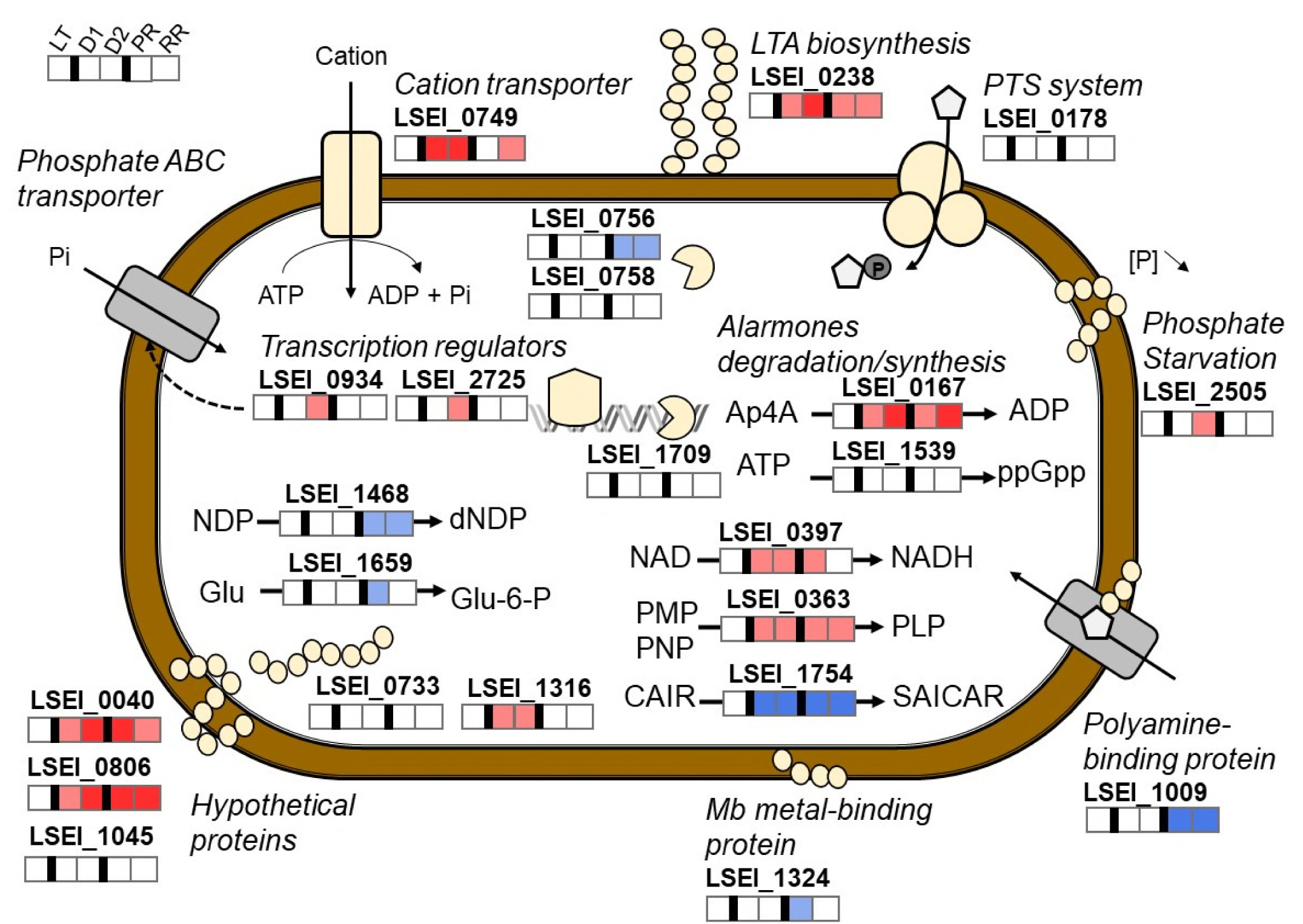
Schematic representation of genes differentially expressed in *L. paracasei* during desiccation and rehydration. Gene expressions are represented by color boxes in light red for upregulation, dark red for upregulations > 2.0, light blue for downregulation, dark blue for downregulation < −2.0 and in white for constitutive regulation. For each gene, boxes from left to right correspond to: lactose incubation (LT), desiccation for 1h (D1), desiccation for 2h (D2), rapid rehydration (RR) and progressive rehydration (PR).

Three genes, LSEI_0934 (a putative DNA binding response regulator), LSEI_2505 (a putative phosphate starvation inducible protein) and LSEI_2725 (a putative sorbitol operon transcription regulator) were upregulated after only 2 h of desiccation and represent potential dry state biomarkers. LSEI_1316 encoding a putative hypothetical protein was upregulated after 1 h and 2 h of desiccation. Interestingly, no gene was specifically upregulated during subsequent rehydration. Conversely, five genes were downregulated specifically during rehydration: LSEI_0756 (a putative hydrolase) LSEI_1009 (a putative spermidine/putrescine-binding protein), LSEI_1468 (a putative ribonucleotide reductase), LSEI_1324 (a putative membrane metal-binding protein) and LSEI_1659 (a putative glucokinase).

## Discussion

Twenty-four desiccation-sensitive mutants were identified from the *L. paracasei* mutant library screening after desiccation followed by rapid rehydration. Contrary to our recent screening of this library for mild stress sensitivity (33), no mutants for putative promoters were identified as sensitive to desiccation. Half of the identified genes were specific of the *L. casei/paracasei/rhamnosus* group. On the contrary, eight genes were well conserved among Gram-positive bacteria. Interestingly, these genes, although present in *Listeria monocytogenes* genome, were not identified by the screening of *L. monocytogenes* random mutants for desiccation resistance (Hingston et al., 2015). Otherwise, this study of *L. monocytogenes* desiccation resistance shows the involvement of a gene encoding a putative glutathione peroxidase, known to prevent oxidative damages. This gene is present in the *L. paracasei* mutant library but was not identified as sensitive. These differences may result from the drying conditions applied such as the surface used for desiccation, the relative humidity, the drying medium or the rehydration conditions.

### Metabolism and transport

Three transporters are required for survival to hydric fluctuation including two putative phosphotransferase systems (LSEI_0178 and LSEI_A05) and one cation transporter (LSEI_0749). Whereas the function of LSEI_0178 is obvious, the LSEI_A05 function as a lactose transporter is uncertain because the corresponding operon is plasmidic and incomplete in this strain. However, this strain is able to metabolize lactose, suggesting that this sugar can enter the cell via another transporter (37). Several authors have reported a link between PTS systems and stress response for *Lactobacillus* genus. *L. plantarum* mutants with an impaired expression of the mannose PTS operon exhibited increased sensitivity to peroxide, probably due to a diminution of glucose capture and energy production (38).

Next, we identified a membrane associated protein (LSEI_1009) predicted to bind polyamines before their transport into the cell by the ABC transporters encoded by other genes of the LSEI_1005-1009 operon. Polyamines have various important physiological roles during stress response, including the modulation of gene expression, signal transduction, oxidative defense mechanism, and cell-to-cell communication (39, 40).

One identified gene encoded a glucokinase (LSEI_1659), a key enzyme of glucose metabolism. In *S. aureus*, glucokinase is also involved in pathogenicity (biofilm formation, virulence factors, cell wall synthesis) (41). In our case, glucokinase as well as PTS interruption could lead to energy diminution and then sensitivity to hydric fluctuations. Another gene, LSEI_0363, encodes a putative enzyme that catalyzes the oxidation of pyridoxamine-5-P (PMP) and pyridoxine-5-P (PNP) to pyridoxal-5-P (PLP) in the vitamin B6 pathway. Vitamin B6, an antioxidant molecule, has been implicated in defense against cellular oxidative stress in *Saccharomyces cerevisiae* (42).

### Cell wall function and structure

Modification at the cell wall level appears to be essential for surviving hydric fluctuation, considering the 6 genes related to the cell wall and identified as essential for hydric fluctuation survival. The gene LSEI_0238 encodes a polysaccharide transporter involved in the export of lipoteichoic acid (LTA), a major constituent of the Gram-positive cell wall (24). Interestingly, this gene was also needed for *L. paracasei* gut establishment (29). The mutant for LSEI_2546, another polysaccharide transporter involved in LTA export, was not identified as sensitive.

LSEI_2505 encodes a putative membrane protein involved in response to phosphate starvation. Although the regulation of this family of genes has been deciphered in *Bacillus subtilis*, their exact role remains unclear (43). Then, LSEI_1324 encodes a putative membrane metal binding protein. Finally, three hypothetical proteins exhibited a transmembrane domain (LSEI_0040, LSEI_0806 and LSEI_1316). Interestingly LSEI_0040 was involved in the adaptation of *L. paracasei* to thermal, ethanol and oxidative stresses (33) and LSEI_0806 to oxidative stress. Both the latter gene and LSEI_1316 were also required for gut establishment (29). The present and previous results show that membrane proteins are important for *L. paracasei* adaptation to various environments and involved in general stress response.

### Regulation of stress response

Two genes involved in alarmone synthesis or degradation are needed for *L. paracasei* survival during hydric fluctuations. Alarmones are putative chemical messengers produced during environmental changes. LSEI_0167 encodes a putative diadenosine tetraphosphatase, an enzyme that hydrolyzes diadenosine tetraphosphate (Ap4A) into two molecules of adenosine diphosphate (ADP). Ap4A has been reported to be involved in heat stress response in *E.coli* (44). Moreover *apaH* (diadenosine tetraphosphatase) mutation causes Ap4A accumulation and sensitivity to thermal stress (45). Nevertheless, Despotović and collaborators wondered whether Ap4A was an alarmone or a damage metabolite because they observed that no signaling cascade was triggered by Ap4A and that its accumulation at high level was toxic for *E. coli*.(46). The second gene (LSEI_1539, *relA*) encoded a putative ppGpp synthetase. The second ppGpp synthetase (LSEI_0901, RelQ) was not present in the mutant library. ppGpp functions as a chemical messenger for metabolism, growth, stress tolerance and virulence in bacteria (47). It is well known that ppGpp interacts with RNA polymerase to affect gene repression or induction. Broadbent et al., (2010) suggested that ppGpp is a signal for acid tolerance response in *L. paracasei*. The LSEI_1539 mutant was also necessary for gut establishment (29).

LSEI_2725 (*gutR*) has been identified as a determinant for surviving hydric fluctuations. Although it is annotated as a sorbitol operon transcription regulator, its predicted function should be reconsidered. Alcantara *et al.,* 2008 reported that in *L. paracasei ATCC334, gutR* is split into 3 ORFs (LSEI_2728, LSEI_2726 and LSEI_2725) as the result of transposition events, and consequently this strain is unable to use sorbitol, contrary to *L. casei* BL23. In addition, the LSEI_ 2724 (*gutM*) gene, encoding a sorbitol operon activator with *gutR*, is dispensable for desiccation resistance. In conclusion, we assume that LSEI_2725 functions as a regulator but independently of the sorbitol operon.

### DNA related enzymes

It was widely reported that desiccation induces DNA damage (1, 50). Indeed, the mutant library screening identified two genes involved in nucleotide metabolism, LSEI_1468 (a ribonucleotide reductase, RNR) and LSEI_1754 (a SAICAR synthase). The RNR mutant was particularly sensitive (mean viability 10-fold less than WT). RNR is an essential enzyme since its mediates the synthesis of deoxyribonucleotides, the precursors of DNA synthesis (51), and we have previously reported that the RNR gene was involved in general response to mild stresses in *L. paracasei* (33). SAICAR synthase is involved in de novo purine biosynthesis (52). These two genes could be involved in DNA repair. Recently, García-Fontana *et al.* (2016) reported that the DNA molecule was overproduced and acted as a protein protector in desiccation tolerant bacteria. This phenomenon could occur in *L. paracasei*. This assumption is supported by the identification of a mutant in a DNA polymerase gene (LSEI_1709).

### Environment sensing

A DNA binding response regulator (RR, LSEI_0934) was required for *L. paracasei* survival of desiccation. LSEI_0934 forms a two-component system (TCS) with histidine kinase (HK, LSEI_0935). The involvement of TCS in *Lactobacillus* response to acid and bile stresses has been reported (54, 55) and the *L. paracasei* ATCC334 genome contains 17 putative TCS (56). LSEI_0934-0935 TCS is organized in a putative operon with five genes encoding phosphate ABC transporters, probably involved in phosphate assimilation. Intracellular poly-P accumulation synthetized from Pi by polyphosphate kinase (Ppk) is a characteristic of some *Lactobacillus* strains to adapt to stress (57). This could be the case during desiccation stress in combination with the phosphate starvation protein gene (LSEI_2505). This assumption should be modulated because mutants for the HK (LSEI_0935) and a phosphate ABC transporter permease (LSEI_0938) are available in the library but were not identified as sensitive to hydric fluctuations.

### Taking into account desiccation and rehydration kinetics

The results indicate that rapid rehydration of WT strain cells was more detrimental than progressive rehydration. It has been reported that progressive rehydration could reduce the stress applied to bacteria as it promotes membrane integrity recovery (58, 59). We found only seven sensitive mutants whatever their rehydration kinetics (Table 4).

Among the genes essential for survival, five genes were expressed at the same level as the control whatever the hydration phase (Table 5). This result again shows the strength of global reverse genetics, since some genes may be essential for a function, although expressed constitutively. Four transcriptomic profiles were observed during hydric fluctuations (Table 5): seven genes were upregulated during both desiccation and rehydration, four genes were upregulated only during the desiccation stage, one gene was downregulated during both desiccation and rehydration, five genes were downregulated only during the rehydration stage. We assume that adaption to hydric fluctuations occurs during dehydration and continues or not during rehydration.

In conclusion, this work identified 24 genetic determinants for the resistance of *L. paracasei* to desiccation. Transcriptomic analysis of the corresponding genes highlighted that seven genes were upregulated during both desiccation and rehydration and four during desiccation only. This analysis provides clues for developing genetic biomarkers to monitor the intensity of desiccation. These findings provide novel insights into the genetic mechanisms involved in desiccation and rehydration adaptation in *L. paracasei*.

## Materials and Methods

### Strains and growth conditions

Wild-type (WT) *L. paracasei* ATCC 334 (CIP 107868, Institut Pasteur Collection) and its corresponding mutants obtained by random transposon insertion mutagenesis (31) were grown statically at 37°C in MRS (Difco). 5 µg/mL erythromycin (Em) was used to select mutants. The mutants correspond to the 1287 genic and intergenic mutants already described (33) (29). For each mutant, the putative inactivated function was assigned thanks to the genome annotation of this strain (24). The mutant library was grown in 96-well plates (200 µl) for 48 h. Plates were mixed (30 s, 700 rpm, Eppendorf MixMate) and 10 µL of each well was used to inoculate 190 µL of MRS in new 96-well plates. Individual mutants were grown in tubes for 48h (2 ml) and vortexed to inoculate at a dilution of 1/100 in new tubes. Plates and tubes were incubated for 24h at 37°C to obtain cells in stationary phase (concentration between 1.0×10^9^ and 2.0×10^9^ UFC/mL). Cells were rehydrated with bromocresol purple (BCP) medium which contains 5 g/L of tryptone, pepsic peptone, yeast extract and sodium acetate, 2 g/L of ammonium citrate and dipotassium phosphate, 1g/L of glucose, 1 ml of tween 80, 0.20 g/L of magnesium sulfate, 0.17 g/L of bromocresol purple and 0.05 g/L of manganese sulfate.

### Drying conditions

The drying chambers were hermetic plastic boxes (20 cm × 13 cm × 6 cm) containing 100 mL saturated potassium acetate (Sigma–Aldrich) solution to obtain 25% RH at 25°C. Samples were placed on a rack in the drying chamber to keep them above the salt solution and the atmosphere was maintained using a ventilator (Sunon, Radiospare, France). Temperature and RH were controlled using an EASY Log USB tool (Lascar Electronics). Drying was performed in a U bottom 96-well microplate (Evergreen, untreated) for mutant library screening and on sterile polypropylene (PP) coupon of 15 mm × 10 mm × 2 mm (Scientix, Fougères, France) for individual treatments. Drying solutions with or without protector were used and as a function of the experiments.

### Screening of the mutant library for viability after desiccation

Stationary growth phase cultures were mixed (1 min, 1000 rpm) and 50 µL of cells were spotted in a U bottom 96-well microplate and centrifuged (5 min, 4000g, 25°C). Pellets were suspended with 50 µL of 50 g/L lactose (1 min, 1700 rpm) and incubated 15 min at 20°C. Then, only 10 µL of suspension were placed in each well and the plates were placed in the ventilated chamber. After 24 h, each well was rehydrated rapidly with 110 µL of (BCP) medium at 37°C. This dedicated culture medium with pH indicator turns to yellow when bacteria metabolize sugars after rehydration. Absorbance at 420 nm (A_420nm_) was measured in a plate reader for 7h at 37°C (Paradigm, Beckman Coulter). Mutants were considered as potentially sensitive when absorbance at 7 h was lower than 1.75, which is the mean value of the WT minus twice the standard derivation, for the two biological replicates. Mutants were selected as potentially resistant when absorbance at 7h (A_7h)_ was higher than 2.07, which is the mean value of the WT plus twice the standard derivation, for two biological replicates. Mutant phenotypes were confirmed, determining viability on plate counts (3 biological replicates).

### Determination of the viability of selected mutants after progressive and rapid rehydration

*L. paracasei* cultures in stationary phase (1 mL) were centrifuged (5 min, 4000 g, 25°C) and pellets were suspended by vortexing with one volume of drying solution composed of lactose 50 g/L. After incubation for 15 min, 10 µL of cell suspensions were placed onto a sterile polypropylene (PP) coupon of 15 mm × 10 mm × 2 mm (Scientix, Fougères, France). Three coupons were prepared for each mutant. The inoculated coupons were placed in the ventilated chamber in plastic petri dishes. For rapid rehydration, 110 µL of BCP medium was deposited on the dried cells and the latter were resuspended by 15 successive cycles using a micropipette. For progressive rehydration, dried cells were introduced into a hermetic chamber at 99% RH for 2 h at 25°C. Then, 110 µL of BCP medium was deposited on the wet bacteria cells that were recovered by 15 successive cycles using a micropipette. Colony enumeration by plate counts was averaged and the viability percentage was obtained with the ratio of colony enumeration (in CFU/coupon) before desiccation to that after desiccation. Mutants exhibiting a viability percentage significantly lower (or higher) than the WT were determined as sensitive (or resistant) (Student test, p < 0.05).

### Bioinformatics

The putative operon organization of corresponding genes was established using the Biocyc website. Then, genes were aligned using BLAST (https://blast.ncbi.nlm.nih.gov/Blast.cgi) against all bacteria to determine their specificity.

### RT-qPCR

Cells in mid-exponential growth phase (OD_600_ between 0.5 and 0.6) were centrifuged and concentrated to an OD_600_ of 20. Cell pellets were suspended with phosphate buffer (10 mM pH 6.5) supplemented with 50 g/L lactose except for control. The suspension was incubated for 15 min at 25°C and 1 mL was deposited on a 0.22 µm polyvinylidene membrane in a glass Petri dish to prevent cell adhesion. Cells were dried in ventilated chambers for 2 h and then subjected to either rapid or progressive rehydration. Rehydrated cells were subsequently detached from the membrane using a cell lifter. Total RNA isolation, cDNA synthesis and qPCR were performed as previously described (60) using TRI Reagent (Sigma Aldrich), DNase I (Roche), iScript™ Reverse Transcription Supermix (Bio-Rad) and SsoAdvanced™ Universal SYBR^®^ Green Supermix (Bio-Rad). Primers were designed by using Primer3Plus (61) (Table 6). Quantitative PCR were performed using a CFX96 Touch™ Real-Time PCR Detection System (Bio-Rad) in triplicate, in a 20 µL-reaction mixture. Cq (threshold value) calculation was determined by a regression model of the CFX Manager™ Software. The relative transcript levels of genes were calculated using the 2^-ΔΔCT^ method (62). In order to select appropriate reference genes, 10 potential housekeeping genes (*fusA, ileS, lepA, leuS, mutL, pcrA, pyrG, recA, recG* and *rpoB*) (63) were tested with all the experimental conditions and analyzed using the CFX Manager™ Software. The genes *fusA*, *lepA* and *rpoB* were selected as the references because they displayed the lowest M values (0.22) and coefficients of variation (0.09), meaning that they have the most stable expression in the tested conditions.

**Table 6.**
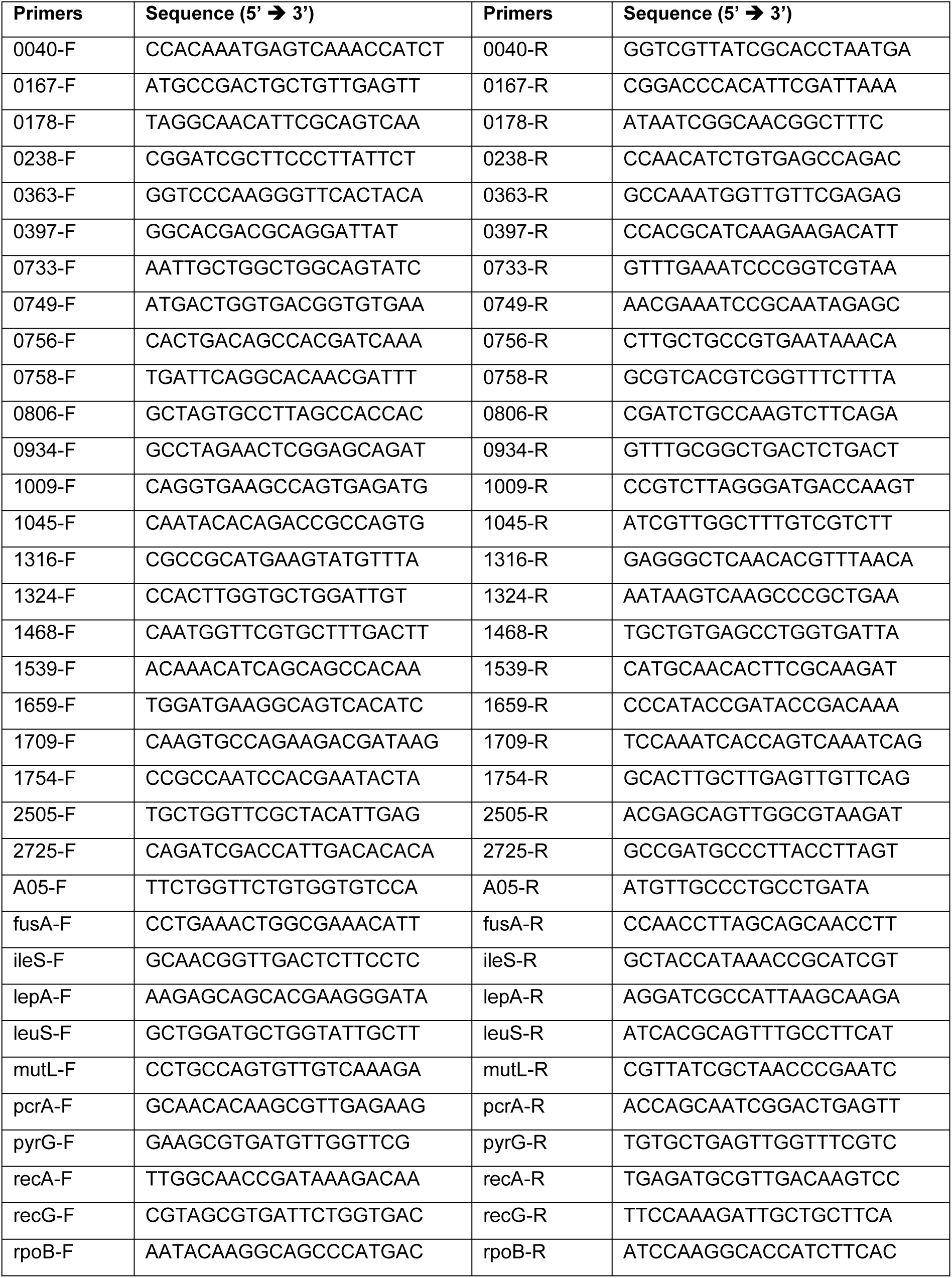
Primers designed for RT-qPCR.

## Author Contributions

Study conception and design: AP, HL, LB and JFC. Acquisition of data: AP. Analysis and interpretation of data: AP, HL, LB and JFC. Drafting of manuscript: AP. Critical revision: AP, HL, LB and JFC. All authors read and approved the final manuscript.

## Acknowledgments

This work was funded by the Regional Council of Bourgogne Franche-Comté and the Fonds Européen de DEveloppement Régional (FEDER). AP was supported by a PhD fellowship from the Ministère de lʼEnseignement Supérieur, de la Recherche et de lʼInnovation (MESRI). We thank the other members of the laboratory for their assistance, including, in particular, Christine Rojas for her skillful technical assistance. We thank the Pasteur Institute Microorganism Collection for the *L. paracasei* strain (CIP 107868).

